# A Deep Dive into Statistical Modeling of RNA Splicing QTLs Reveals New Variants that Explain Neurodegenerative Disease

**DOI:** 10.1101/2024.09.01.610696

**Authors:** David Wang, Matthew R. Gazzara, San Jewell, Benjamin Wales-McGrath, Christopher D. Brown, Peter S. Choi, Yoseph Barash

**Author notes:** Passed away March 18th, 2023 (Christopher D. Brown).

## Abstract

Genome-wide association studies (GWAS) have identified thousands of putative disease causing variants with unknown regulatory effects. Efforts to connect these variants with splicing quantitative trait loci (sQTLs) have provided functional insights, yet sQTLs reported by existing methods cannot explain many GWAS signals. We show current sQTL modeling approaches can be improved by considering alternative splicing representation, model calibration, and covariate integration. We then introduce MAJIQTL, a new pipeline for sQTL discovery. MAJIQTL includes two new statistical methods: a weighted multiple testing approach for sGene discovery and a model for sQTL effect size inference to improve variant prioritization. By applying MAJIQTL to GTEx, we find significantly more sGenes harboring sQTLs with functional significance. Notably, our analysis implicates the novel variant rs582283 in Alzheimer’s disease. Using antisense oligonucleotides, we validate this variant’s effect by blocking the implicated YBX3 binding site, leading to exon skipping in the gene MS4A3.

## Introduction

Genome-wide association studies (GWAS) have implicated thousands of genetic variants in human complex traits and disease. However, elucidating the functional mechanisms through which a variant acts on a trait remains challenging. The current prevailing hypothesis maintains that the effect of a variant on a trait is mediated by its effect on gene expression either by directly affecting trans acting elements like transcription factors or by disrupting cis regulatory elements such as transcription factor binding sites. This view is supported by the observation that GWAS associations are enriched for eQTLs in non-coding regions, leading to many studies reporting a high degree of colocalization between eQTLs and GWAS signals [1]. However, many GWAS hits still cannot be explained by eQTLs thus many alternative hypotheses have been proposed to potentially address this gap [2]. For example, several studies have suggested that eQTLs are context specific [3] while others claim that QTLs for other molecular traits could be more informative. Specifically, a comprehensive study of many molecular QTLs identified RNA splicing QTLs (sQTLs) as a key contributor to several disease phenotypes [4]. However, sQTLs that explain GWAS variants are still comparatively underrepresented in recent studies. We postulate that improved sQTL detection, and consequent overlap with GWAS signals, can be achieved by focusing on three main elements.

First, the RNA splicing representation used for sQTL discovery should ideally capture the full complexity of splicing variations in the transcriptome. Splicing can be accurately measured using short read RNA sequencing and quantified at the local “event” (e.g. cassette exon) or junction level using tools like MAJIQ [5, 6], Leafcutter [7] or rMATs [8]. It can also be quantified at the isoform level with transcript expression using tools like Salmon[9], Kalisto [10] or RSEM[11]. However, each splicing quantification approach presents only a partial view of splicing variation and joint analysis of event and isoform representations can yield complementary insight. For example, event based methods such as rMATs miss denovo junctions and exons, while methods such as Leafcutter miss intron retention. All event based methods miss alternative transcript starts and ends which can be captured by isoform based approaches. However, isoform quantification typically relies on annotations and can’t handle unannotated transcripts [12]. Usage of long reads can greatly improve unannotated transcript detection and quantification but low coverage and high-error rates remain a challenge [13].

Second, statistical models for sQTL effect size inference could be improved by accounting for the discrete and heteroskedastic nature of RNA splicing data. Most modern sQTL analysis frameworks can be traced back to eQTL mapping. As such, the methods combine linear regression with various phenotype normalization procedures such as the rank inverse normal transform (RINT). However, RNA splicing quantifications are quite different from gene expression estimates. Splicing is typically quantified as percent splice-in (Ψ) which is a measure in the domain of [0,1] of relative junction or isoform usage. Moreover, Ψ values are derived from sparse discrete read counts that span splice junctions. Accounting for the consequent uncertainty in these measurements by modeling read counts can likely increase detection power and better control false discoveries. This is especially important in contexts where variant associations with splicing are confounded by associations with expression and coverage which may introduce heteroskedastic effects. Furthermore, splicing is a multivariate phenotype with most genes containing multiple junctions which requires additional considerations in statistical model design and multiple hypothesis testing correction.

Finally, there are several covariates that affect the power of marginal SNP-junction associations which should be considered to maximize discovery power. One covariate is the proximity of a SNP to a splice site. Most variants that are associated with splicing are near splice sites and disrupt core spliceosome or RBP binding motifs [14]. Another covariate is the coverage of the junction. Junction coverage can be affected by the sequencing depth of experiments, the expression of the gene, and proximity to the 3’ end. The power to detect sQTLs is greatly reduced at low coverage since the variance of statistical estimators is higher and effective sample size is smaller due to dilution caused by imputation of missing values. Currently, sQTL analysis pipelines typically handle these covariates implicitly by only considering SNPs within a fixed window around a gene’s transcription start site and filtering events beyond a certain level of missing values. However, detection power could be further improved by allocating the multiple hypothesis testing budget in a way that favors sQTLs which are more likely to be detected.

Previous works have attempted to improve various aspects of sQTL modeling. Many studies acknowledge that unannotated (denovo) splice junctions are important and therefore use Leafcutter [7] for detecting such junctions. Indeed, our analysis [6] shows LeafCutter offers an efficient method for detecting splicing variations, but it is not able to capture intron retention or alternative starts and ends. Other studies consider the count-based nature of splicing using generalized linear models like GLiMMPS [15] or DRIMSeq [16] but do not account for multiple splice junctions per gene. On the other hand, methods like sQTLSeeker [17] and THISLE [18] account for the multivariate nature of splicing phenotypes through multivariate testing or tests for heterogeneity respectively, but do not model the limited count data used in RNA splicing quantification. Finally, approaches like Independent Hypothesis Weighting (IHW) [19] have been proposed to handle covariates associated with power but are designed to control false discovery rates (FDR) for independent tests. In contrast, the highly correlated nature of splicing events and SNPs in sQTL analysis requires relaxed independence assumptions [20].

In this work, we first investigate the potential limitations of existing sQTL methodologies. Based on our findings, we then propose MAJIQTL, a new statistical framework for improved sQTL discovery and analysis. Our pipeline includes a comprehensive splicing phenotype representation derived from MAJIQ quantifications, a weighted multiple hypothesis testing correction method for sGene discovery utilizing covariates, and a Beta-Binomial composite testing model for accurate and interpretable sQTL effect size inference. These components not only find more sGenes/sQTLs, but also effectively prioritize the variants with functional significance. We demonstrate the validity and utility of our approach through extensive simulations, real data applications, and assessment of variants’ effect by orthogonal methods such as splicing prediction algorithms (splicing codes) and molecular experiments. When applied to the GTEx dataset, our method finds significantly more functionally relevant sQTLs which are enriched for important functional annotations and co-localize with neurodegenerative disease GWAS variants. Notably, we find and validate rs582283, a variant that is implicated in Alzheimer’s disease. We show this variant disrupts the RBP binding site of YBX3 in the gene MS4A3 leading to skipping of exon 7, providing a plausible mechanism of action. We make our results easily accessible through a dedicated web-tool which can be further explored with VIOLA-QTL, a new sQTL visualization tool. Taken together, our results illustrate the power of careful examination and consequent improvement of multiple aspects of sQTL analysis methodology, leading to novel sQTLs which explain additional GWAS signals.

## Results

### Analysis of GTEx Tissues Points to Potential Improvement for sQTL Modeling

We start this study by first applying a standard sQTL (denoted std-sQTL) pipeline to samples from five representative tissues in the GTEx dataset: whole blood, lung, brain - cerebellum, liver and heart (see Supplementary Note for details). This pipeline was popularized by the GTEx consortium [21] and has since been adapted by dozens of modern studies with great success. We decomposed the std-sQTL pipeline into three elements: input splicing representation, statistical models for effect size inference and power informative covariates. We then evaluated each of the three elements and investigated how it could be potentially improved. Before we turn to describe this investigation we note that although we focus on these three elements, we also identified other important areas for consideration in the std-sQTL pipeline such as mapping, filtering and confounder correction. To maintain scope, we defer those to the Supplementary Note (Figure S1).

In the first step of sQTL detection assessment, we used the std-sQTL pipeline to call sGenes and assessed the effect of using different splicing representations. For a splice junction/event level representation, we quantified splicing (Ψ) using Leafcutter’s intron cluster approach (most common method used in the std-sQTL pipeline) and MAJIQ’s local splice variation (LSV) approach. For an isoform level representation, we quantified transcript expression using Salmon TPM and normalized these values to obtain the percent abundance of isoforms per gene. Although several studies highlighted the limited accuracy of whole transcript quantification from short read RNASeq data, numerous studies have used this approach for transcript QTL (trQTL) [18, 22] so assessing the value of this approach is warranted. The use of MAJIQ LSVs resulted in the discovery of a similar number of sGenes compared to the use of Leafcutter intron clusters across all tissues when only considering annotated (sGenes) and denovo (dGenes) splicing events (Figure 1a). However, the std-SQL pipeline reports between 1.4 times to 2 times more sGenes (or iGenes) with intron retention events when using MAJIQ quantification of intron retention (IR) in heart and cerebellum respectively. These genes are otherwise not significant when only considering splicing events without intron retention (Figure S2). The use of Salmon isoform quantification also identified sGenes (isoGenes) that do not overlap those identified by junction level approaches. These sGene hits are enriched for isoforms that differ at transcript starts or ends. Overall, we find that using MAJIQ with transcript quantifications in the std-SQTL pipeline can capture two fold more genetically associated splicing variations compared to using Leafcutter.

**Figure 1:**
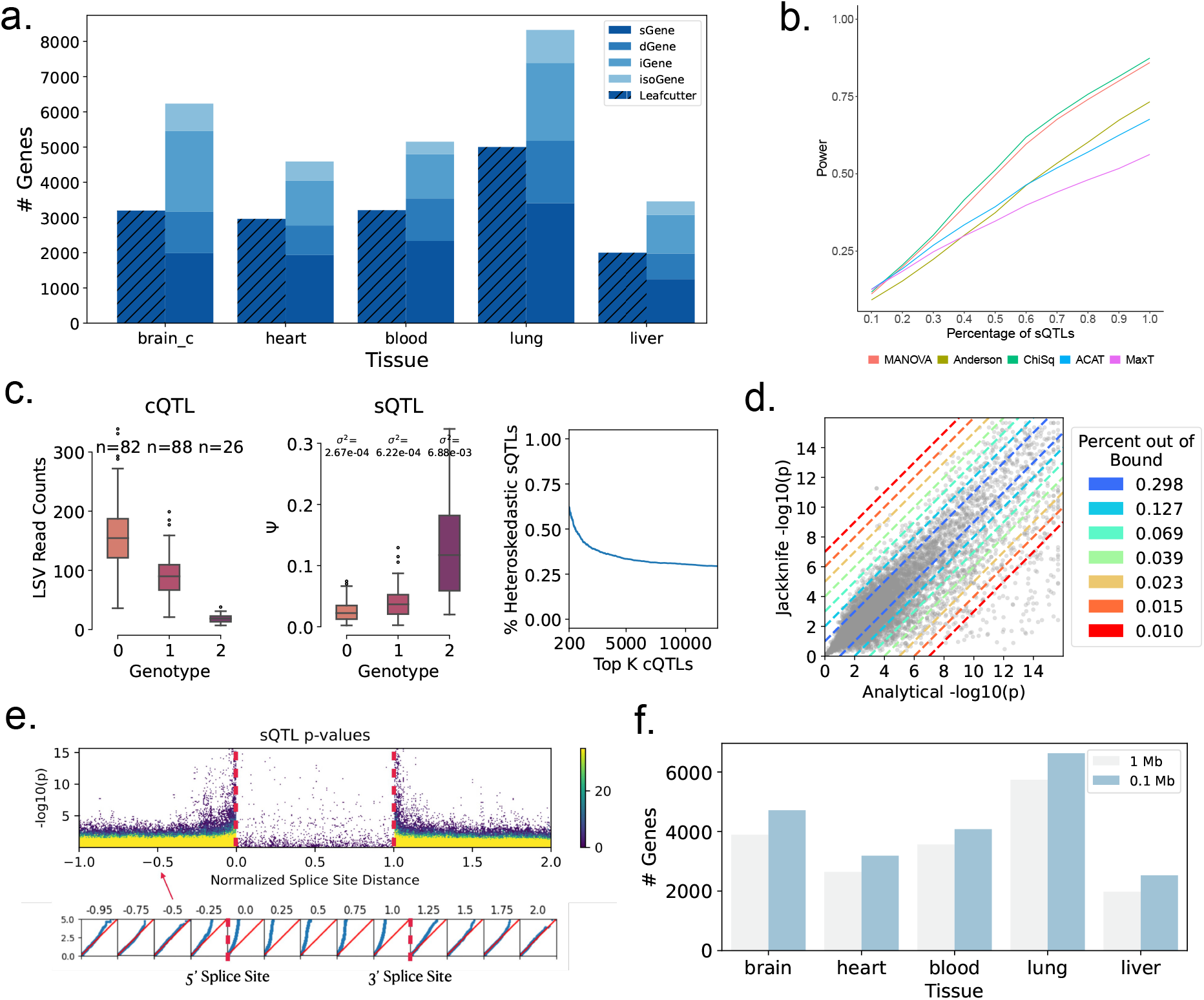
GTEx Case Study. **(A)** Barplot showing the number of sGenes found by MAJIQ and Leafcutter across 5 representative tissues. The unique sGenes are stratified by whether they are called due to annotated events (sGenes), denovo events (dGenes), intron retention (iGenes) or isoforms (isoGenes). Using Leafcutter alone fails to find iGenes and isoGenes. **(B)** A comparison of the statistical methods for sGene discovery. The x-axis represents the percentage of tests in a locus that are sQTLs. For example, 0.2 indicates that there are 2 junctions out 10 where the alternative hypothesis is true. The y-axis shows the power. **(C)** An example of heteroskedasticity introduced by coverage. The left plot shows that this event is a cQTL in which the genotype is correlated with coverage. The middle plot shows that the event is an sQTL in which the genotype is correlated with Ψ. However, the variance is highest when the coverage is low and the variance is lowest when the coverage is high. The right plot shows that the percent of top K cQTLs that are also heteroskedastic sQTLs (based on Bruesch-Page test) increases as K decreases. **(D)** A comparison of jackknife p-values (y-axis) and OLS analytical p-values (x-axis) when applying linear regression to splicing data. Due to model misspecification (i.e. heteroskedasticity), the values do not agree well. The colors indicate bounds which represent minimum fold differences. For example, the bound at 1 (blue) represents at least a 10 fold difference in -log10 p-value between the jackknife and analytical values for all points to the left and right of the bound. The numbers in the legend incidate the percentage of points that fall beyond the bound. **(E)** P-value inflation as a function of the SNP’s distance to the 5’ and 3’ splice sites. The normalized distance is shown for a 1Mb window. The QQ plots below show that the distribution of the observed p-values is far from uniform for sQTLs closer to slice sites. The colors represents the density of data points. **(F)** The number of sGenes discovered in each tissue depending on whether a 1 Mb (blue) or 0.1 Mb (grey) window was used for the std-SQTL pipeline.

Second, we assessed the statistical methods used to detect variants associated with sGenes and sQTLs. The primary challenge with sGenes is the multivariate nature while the challenge with sQTL effect size inference lies in the data. Splicing phenotypes such as those quantified by LeafCutter or MAJIQ are derived from multiple splice junctions in a gene. Thus identifying sGenes requires detecting non-zero effect sizes between a single genetic variant and multiple splice junctions. Various models have been proposed to handle the multivariate and discrete aspects of this phenotype. First, we consider the multivariate nature of the splicing phenotype. There have been several methods proposed for multivariate testing to detect sGenes. This includes MANOVA, Anderson’s pseudo F test implemented in sQTLSeekeR2 [17], sum of *χ*^2^ test implemented in THISLE [18], ACAT[23], and exact FWER control implemented in QTLTools [24] and used in the std-sQTL pipeline. We provide a summary of each method and their mathematical relationships in the Supplementary Notes. To assess how these methods behave on splicing data, we simulate genes with correlated splice junctions and make a subset of the junctions be associated with a genetic variant. The details of this simulation study are given in the Supplementary Notes. Then we evaluate the power of each method to detect sGenes. Figure 1b shows MANOVA and *χ*^2^ perform well compared to sQTLSeekeR2, ACAT and FWER control. However, since it is difficult to generalize MANOVA to non-Gaussian distributions, we do not consider this class of approaches any further. Note that a Dirchlet-Multinomial regression model cannot work as a generalization of MANOVA in this setting since each LSV or intron cluster is a multinomial unit and there are multiple LSVs in each gene. Interestingly, the performance of sQTLSeekeR2 eventually surpasses FWER control when 40% of the junctions in a gene are associated with the variant. Overall, these result point to potential improvement in sGene detection power by using alternatives to the commonly used FWER control.

Next, we assess whether using linear regression with RINT transformations (std-sQTL approach) for effect size inference between single SNP-junction pairs is appropriate given that splicing quantifications are derived from discrete junction spanning read counts. First, we note that Ψ values generally follow a distribution skewed toward 0 or 1 and thus residual estimates are not normal. However, given the population scale data of modern QTL studies, linear regression models used in this context are still robust because the test statistic maintains asymptotic normality at reasonable convergence rates as a consequence of the central limit theorem [25]. Nonetheless, violations of Gauss Markov assumptions, such as heteroskedastic errors, can result in miscalibrated models regardless of sample size (Supple-mentary Note). Indeed, we show that large heteroskedastic effects are persistent in the data due to the interplay between expression and splicing (Figure 1c). Specifically, a variant that affects expression (eQTL) can cause the splice junction coverage (and consequently variance of its quantification) to vary with genotype (cQTL). The example in this figure shows a gene in which the variant affects coverage and Ψ. The variance of Ψ increases from 2.67e-4 to 6.88e-3 as the coverage decreases (Figure 1c left and middle). Furthermore, we show that the percent of top K cQTLs which are also heteroskedastic sQTLs (significant Breusch-Pagan p-value) increases as K decreases (Figure 1c right). The genotype induced differences in coverage can also lead to differences in quantification resolution which effectively renders them discrete. For example, when the total coverage is 10, there are only 11 possible values of Ψ while there are 101 possible values of Ψ when the coverage is 100. This phenomenon cannot be addressed by the commonly used RINT transformation which assumes that the phenotype is homoskedastic and continuous. Thus it becomes necessary to select a generalized linear model (GLM) with a variance function that is properly specified for this data. We compare the goodness of fit of Beta, Binomial and Beta-Binomial models as well as a previously proposed Binomial model with mixed effects [15] against a Gaussian model using residual quantiles [26] on the lead SNP-junction pairs in sGenes (Figure S3). Surprisingly, the binomial model has the worst fit while the beta binomial has the best fit, suggesting that modeling overdispersion is crucial. Furthermore, we compute the agreement between the analytical and jackknife p-values to measure calibration of the linear regression model and to assess false positive control (Figure 1d). We show that 29.8% of sQTLs have a 10 fold difference in p-value (a shift of 1 decimal place) while 1.0% have a 70 fold difference. Interestingly, the misspecified linear model generally has lower analytical p-values compared to jackknife p-values (points mostly below the x = y line) suggesting an inflation of false positives. An important point to make in this context regards the common practice to test against synthetic nulls generated through data permutations in order to assess false positive rates [27]. Such synthetic null generation pools the variance, thus removing the effects of heteroskedasticity which consequently underestimates false positives and is not appropriate here.

Finally, we investigate potential covariates associated with the power of marginal associations. When conducting multiple tests, it is often the case that a researcher may want to make fewer tests based on a covariate. The covariate ideally informs the a priori chance of a significant signal to enable efficient filtering of uninformative hypotheses and reduce the multiple hypothesis testing burden. It is common practice to implement basic filtering based on covariates. The std-sQTL pipeline for example, tests SNPs up to 1 Mb away from a gene’s transcription start site and filters junctions which are missing in more than 50% of samples. Other studies are more conservative, filtering junctions missing in 10% or more of the samples [28]. While sensible, these filters were borrowed from eQTL studies [29] and may be less effective in the sQTL setting. In subsequent analysis, we find that proximity to splice sites (Figure 1e) and the missingness rate of a junction (Figure S4) are two covariates associated with sQTL detection power. The missingness rate is associated with coverage since missing values are more prominent at low coverage and are MNAR (missing not at random) rather than MCAR (missing completely at random) (Figure S5). Using a naive filtering approach, we show that reducing the 1 Mb window around a gene’s TSS to a 0.1 Mb window around each junction’s splice site significantly increases the number of associations (Figure 1f). Furthermore, increasing the number of tested junctions up to a 50% missingness rate greatly increases total discoveries (Figure S6). Notably, such a threshold based filtering approach can be seen as a special case of weighting where the filtering criteria is not correlated with the null test statistic (Supplementary Note). With fixed thresholds, the filtered cases have their weights reduced to 0 and the remaining budget is redistributed uniformly across the remaining tests. However, it has been shown that when correcting for multiple hypothesis testing, a more nuanced approach of redistributing the test budget based on a covariate can improve power [19]. Thus, we conclude from this analysis that while optimal hypothesis weighting by covariates for sQTL testing is not known, it is evident that improvements can be made to increase power.

### MAJIQTL Offers a Robust and Powerful Statistical Framework for sQTL Discovery and Analysis

Motivated by the above observations, we developed MAJIQTL, a fast and light-weight statistical framework for sQTL discovery and analysis that builds on the popular MAJIQ method for RNA splicing quantification[5, 6]. Specifically, MAJIQTL features an exhaustive input splicing representation that integrates splice junction and full isoform level information; a weighted multiple testing method which improves sGene detection power by leveraging covariates; and a robust and interpretable regression model for sQTL effect size inference and variant prioritization. MAJIQTL is tightly integrated with the MAJIQ family of tools and contains many updates to those which we expand upon below. We now turn to describe each of the MAJIQTL components.

In the first step of the analysis pipeline (first column in Figure 2), we compile a comprehensive splicing phenotype representation combining splice junction quantifications from MAJIQ and full isoform level quantifications from Salmon. This allows MAJTIQTL to capture complex splicing variations involving unannotated junctions and exons, as well as intron retention and alternate transcript starts/ends. Integration with MAJIQ also enables use of its builtin event type classifier, which can annotate both classical and complex events into interpretable units (exon skipping, Alt 3’, Alt 5’, etc), and MOCCASSIN, a dedicated tool for splicing confounder correction. Furthermore, we introduce various best practice guidelines for splicing phenotype processing such as filtering procedures (Supplementary Note).

**Figure 2:**
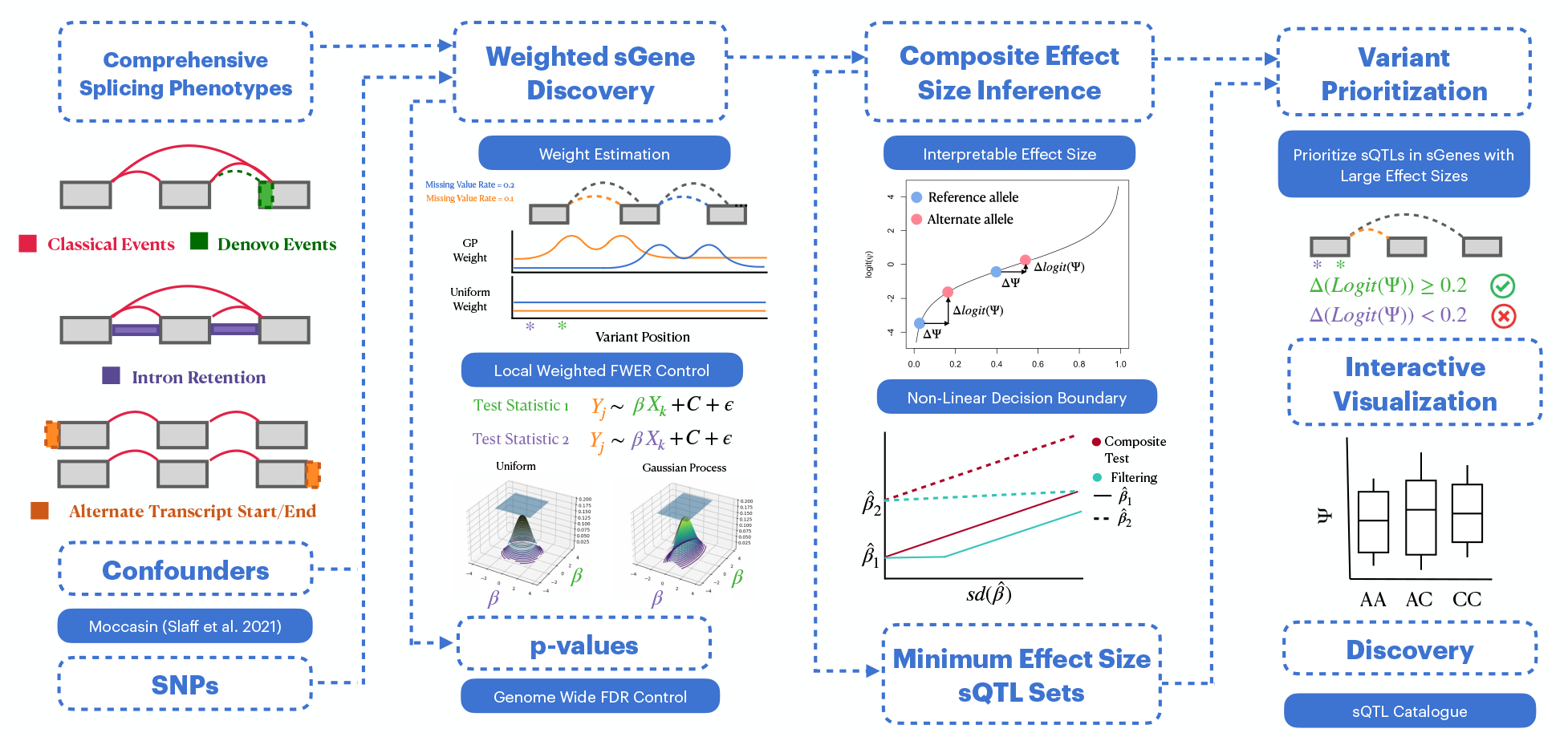
MAJIQTL Pipeline. An overview of the MAJIQTL pipeline. First (column 1), we quantify splicing using MA-JIQ and combine these quantifications with isoform ratios, creating a comprehensive set of splicing events which includes classical, denovo, intron retention and alternate transcript start/ends events. Confounding effects are corrected using MOCCASIN. Second (column 2), we use our weighted multiple testing method to discover sGenes. This approach uses a GP regression model to learn a mapping to weights from covariates based on the missingness rate of the junction (orange/blue) and the proximity of the variant to the splice site (purple/green). The baseline uniform weight model is shown for comparison. Then we use these weights to compute a gene level p-value that controls the FWER of sQTL discoveries in each gene. Intuitively, FWER control is achieved by assigning a threshold to each sQTL’s test statistic (represented by the edges of the blue square) subject to a constant budget constraint (volume of the density plot directly under the square). Under a uniform model, the thresholds are all equal (left). However, our weighting model assigns thresholds proportional the weights (right). Third (column 3), we use a composite Beta-Binomial regression model to estimate effect sizes of sQTLs identified in the previous step. The effect size estimate has a fold change interpretation which is related to the ΔΨ measure of splicing change (x-axis) through the logit function (y-axis). Then we then use a composite testing approach to create sets of sQTLs defined by a minimum effect size. An advantage of this approach (red) is the non-linear decision boundary for assigning sQTLs to the set which accounts for the variance of estimates unlike filtering by observed effect sizes (blue). Finally (column 4), we can use these sets to prioritize sQTLs with large effect sizes. Our results can be visualized using the VOILA-QTL package in MAJIQ and we report a catalogue of our sQTLs in the MAJIQlopedia database.

In the second step of MAJTIQTL’s analysis pipeline (second column in Figure 2), we introduce a new weighted multiple hypothesis testing method to improve sGene discovery power. Accounting for multiple testing is imperative given the large number of tests performed between all pairs of splice junctions and the cis SNPs in a predefined locus around each gene. However, it has been shown that the naive approach of applying a single multiple testing correction procedure (e.g. Benjamini-Hochberg) to all tests fails to control false discoveries [30]. Instead, modern sQTL studies rely on a hierarchical correction strategy [21]. This approach first controls the sGene false discovery rate (FDR) at a desired level (e.g. FDR = 0.05) by using gene level p-values computed through the max T method to learn a decision threshold *α*. By rejecting only sGenes with a p-value below *α*, the family-wise error rate (FWER) of all tests in each gene is controlled at level *α*. This approach is able to properly handle the complex local correlation structure and number of tests unique to each gene unlike the naive approach. However, rejecting the max T p-value at level *α* represents unweighted FWER control where each test is valued equally in the FWER budget (i.e. *α*). Our case study shows that sQTLs are more likely to appear near splice sites or in junctions with low missingness rate. Thus we instead want the p-value to reflect weighted FWER where the FWER budget is allocated in manner that is proportional to the power informative covariate.

To address this need, MAJIQTL computes a weighted p-value for each sGene that increases discovery power while controlling local FWER under the null. The inputs are the covariates and test statistics for each sQTL. Given these inputs, the method first uses an empirical Bayes approach to learn a flexible non-parametric function that maps the covariates to weights that are proportional to the posterior probability of the corresponding test being alternative *P* (*H*_1_) using a Guassian Process (GP) regression (Figure 2). Notably, the GP function is optimized jointly over all exons, genes, and SNPs to avoid over-fitting and information leakage. Next, our MAJIQTL computes a gene level p-value using these weights such that rejection of the p-value at level *α* maintains a constant FWER budget but redistributes the budget proportional to the weight of each test (bottom of second column in Figure 2). The details and theoretical guarantees are fully described in Methods. However, there are two key innovations that we highlight here. First, the estimation of *P* (*H*_1_) does not require any numerical optimization. Instead, we use the Kolmogorov-Smirnov distance as a “plug-in” estimator and scale this value by a learned parameter *λ* which maximizes sGene discoveries. We will show that this estimator is a strikingly accurate approximation for the ideal weight *P* (*H*_1_). Second, the null distribution of the weighted test statistic cannot be computed analytically. We will show it can be sampled quickly and accurately from a matrix normal distribution, thus completely avoiding the use of permutation procedures. These two strategies enable our approach to efficiently scale to hundreds of millions of tests and only takes minutes to run on the GTEx datasets.

In MAJIQTL’s third step (third column in Figure 2), we introduce a model for variant prioritization and effect size inference. A widely recognized limitation of current sQTL pipelines is the lack of dedicated methods for selecting candidate variants in sGenes. For example, the GTEx portal only reports a single sQTL (lead SNP) with non-zero effect on splicing in each sGene. However, a researcher may be interested in all the sQTLs with the largest effect sizes (identifying pathogenic splice site disrupting variants) or may want to omit the sQTLs with small effect sizes as those may be hard to validate experimentally and may be less attractive for assessing phenotipic effects. Thus we would ideally like to form sets of variants defined by a minimum effect size threshold such that all variants in the set exert an effect on splicing that exceeds that threshold. We can then prioritize the sets defined by larger thresholds in downstream analysis. However, as shown by our case study, effect size estimates from existing models are limited by misspecifcation and thus not comparable between sQTLs. Specifically, excessive phenotype normalizations obfuscate inference of effect sizes that can be interpreted on the same scale. Furthermore, even when using a well calibrated model, simply filtering by the effect size estimator without considering confidence in the estimate may be inappropriate [31]. Notably, even though effect sizes are considered for eQTL [32], this issue of confidence intervals around effect estimates is still not incorporated in the GTEx and similar eQTL pipelines.

To address these needs for variant prioritization and effect size inference, we propose using a composite Beta-Binomial model to identify thresholded rejection sets. These are illustrated at the bottom of the third column in Figure 2. In brief, this variant prioritization strategy starts with all the sQTLs that pass the FWER bound in each sGene which have confirmed non-zero effect size. Then we use a beta-binomial regression model on this data to infer effect size magnitude 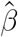(Methods). This allows us to handle uncertainty in splicing quantifications due to coverage and heteroskedastic effects and the coefficient naturally has a fold change interpretation which is the log odds of junction inclusion (Figure 2). This measure for effect size is appropriate as it has been shown that splicing dynamics are a competition for splice site usage and splicing changes induced by variants in *cis* can be modeled well using a sigmoid like switch between splice sites [33]. To define the composite rejection set, rather than rejecting variants under the null hypothesis 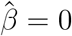, we reject under the composite hypothesis 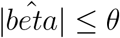 where *θ* is a lower bound on tolerable effect size. This testing procedure, similar to ones previously applied for gene expression changes [34], is described in Methods.

The MAJIQTL methods are deeply integrated into the popular MAJIQ software package for RNA splicing analysis. They are designed as “plug-in” replacements for the commonly used Leafcutter and QTLTools with comparable inputs and outputs. Specifically, MAJIQTL can be run without any programmatic interface, requiring at minimum only a BED file containing splicing quantifications and a VCF file containing variant information to perform all operations. Using these standard files as input, the MAJIQTL software is equipped with several additional tools that improve speed and user accessibility. First, we implemented a new library for the Beta-Binomial model which achieves improved speed and accuracy though parallelization and use of exact gradients in optimization. Second, we include a dedicated interactive visualization tool in VOILA-QTL (bottom right illustration in Figure 2). VOILAQTL is an extension of MAJIQ’s VOILA visualization tool with a slew of new features to enable visual analysis of sQTLs (Figure 2). Finally, our results are distributed through the MAJQTL database which is easily accessible online (see Data Availability). We note that although our approach is designed to integrate and take advantage of MAJIQ’s features, our pipeline is still compatible with other splicing quantification tools such as Leafcutter and rMATs.

### MAJIQTL Weighted Multiple Testing Improves sGene Discovery Power

We first evaluate the power of our weighted multiple testing method to discover sGenes in the five representative GTEx tissues. We obtain a p-value for each gene by providing the model with summary statistics computed using RINT OLS between every splice junction and cis SNP within a 1 Mb window around the gene and matching covariates for each junction and SNP. The two covariates are the normalized distance of a SNP to the paired junction’s splice site and the missingness rate of each splice junction across all samples. We also apply the independent filtering method to these datasets. This baseline method reports the standard max T p-value for each gene for incrementally decreasing cis window sizes and can be interpreted as a binary weighting approach (filtered tests have zero weight and the remaining tests have uniform weight). The results for brain-cerebellum are shown in Figure 3 while the results for the remaining tissues are shown in Supplementary Note. For a given FDR rate, the power of our method is significantly improved compared to the baseline (Figure 3a). At a FDR of 0.05 our method finds 12% more sGenes than independent filtering at a 0.1 Mb window size. This figure also illustrates the behavior of optimizing the scaling parameter which achieves optimal power on this dataset at *λ* = 5.

**Figure 3:**
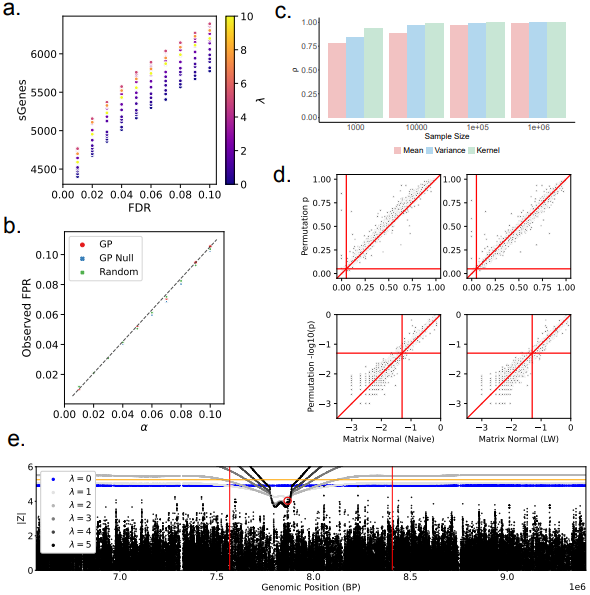
Weighted Multiple Testing Evaluation. **(A)** The number of sGenes (y-axis) discovered by MAJIQTL’s weighted multiple testing method in brain - cerebellum at varying FDR levels (x-axis). For a given FDR level, the color of each point indicates the value of the model parameter *λ* which was optimized to maximize sGene discoveries (*λ* = 5). When *λ* = 0, the behavior of the method is equivalent to the unweighted max T sGene discovery method used by GTEx (baseline). **(B)** The observed false positive rate (y-axis) of the method at varying p-value cutoffs *α* (x-axis). The method was applied to synthetic null data generated using permutations and the observed FPR matches the expected FPR (x=y line). The colors represent 3 different approaches used to select the weights: the GP trained on the original data (red), the GP trained on the null data (blue), and random weights drawn from a uniform distribution between 0 and 1 (green). **(C)** The Spearman correlation *ρ* between *P* (*H*_1_ |*C*) and the pooled KS statistic estimator 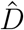 at varying sample sizes (the number of sQTL summary statistics used to estimate the pooled KS statistic). The colors represent 3 different models for the alternate distribution. Under the mean (red) and variance (blue) models, the mean and variance of the alternate distribution vary with *P* (*H*_1_|*C*) respectively. The kernel model (green) uses the empirical kernel density of real sQTL summary statistics to approximate an alternate distribution. **(D)** The correlation between the unweighted max T p-values where the null distribution is computed using permutations (y-axis) or our matrix normal (MN) sampling approach (x-axis). The left panels show results using the naive MN sampling approach (sample covariance estimator) while the right panels show results using the Ledoit-Wolf shrinkage estimator. The top panels show the raw p-values while the bottom panels show the -log10 p-values. **(E)** An example of the weighted FWER bound computed by the method for the locus around a gene with position shown by vertical red lines. For clarity, only the bound for a single junction is shown. The FWER of all tests in the locus is controlled at the BH critical value (0.022) which controls gene level FDR at 0.05. The greyscale colors indicate the value of *λ* for which each bound was computed. Recall that when *λ* = 0, the method is equivalent to the unweighted max T method and produces a uniform bound (blue). For comparison, the Bonferroni bound is shown in orange. This gene is considered an sGene using the weighted method since it contains at least 1 sQTL with a test statistic (y-axis) above the bound (red circle).

Next we confirm that our method still maintains false positive control under the null. To show this, we first generate synthetic null datasets using a permutation scheme that preserves the correlation between junctions and between SNPs (Supplementary Note). Then, we show that our method controls false positives by producing p-values that are uniform when applied to this null data (Figure 3b). It is reasonable to assume that false positive control is only achieved since the weights estimated by the GP trained on null data are approximately uniform and thus the weighted p-value is similar to the standard max T p-value. However, we find that our model’s p-values are well calibrated regardless of whether we trained the GP on the null data or original data (non uniform weights). Furthermore, we observe the same degree of false positive control using weights randomly drawn from a standard normal distribution. These result together point to the robustness of our approach in achieving control regardless of how the weights are estimated.

The power of our method depends on how well the GP regression model is able to map covariates to *P* (*H*_1_ |*C*). However, as previously described, we instead use a strategy of mapping to a plug-in estimator (KS distance between the null and observed empirical distribution of test statistics) and then scale this value to maximize sGene discoveries. Here, we explore how well this approach approximates a mapping to *P* (*H*_1_|*C*). We perform a simulation where we draw test statistics from a two component mixture of normal distributions where each component represents the null and alternative distributions. The mixing proportion, *P* (*H*_1_|*C*), is assumed to be linearly correlated to the covariate. We note that our model can theoretically handle a non-monotonic relationship between *P* (*H*_1_|*C*) and the covariate, however such behavior is not observed in real sQTL datasets and we therefore do assess such relations. The full details of the simulation are given in Supplementary Note. Using the aforementioned simulated data we then show that *P* (*H*_1_|*C*) can be approximately recov-ered using our modeling approach. For this,, we use Spearman correlation to measure the agreement between the ground truth values of *P* (*H*_1_|*C*) and the KS statistics, as shown in Figure 3c. We used three different settings for the simulated data: Data were the mean (red bars) or variance (blue bars) of the alternative distribution was correlated to *P* (*H*_1_|*C*), or data generated such that the distribution was empirically constructed from real sQTL data (green bars). These results illustrate that our fast approximation for the weights results in near optimal power.

Next, we turn to show the quality of approximating the null distribution through sampling. The test statistics for a gene are assumed to follow a matrix normal distribution with mean zero under the null. We can sample from this distribution instead of using permutations to generate an empirical null, but the quality of this approximation is limited by the estimation of the covariance matrices. In brief, the covariance matrices can be naively estimated using the sample correlation between SNPs and between junctions. This approach has been used in similar methods like gene-MVN for eQTLs [35]. However, when the number of features (SNPs) greatly exceeds the number of samples, the quality of the covariance estimator is reduced. We remedy this using the Ledoit-Wolf shinkage estimator. We compare the naive sampling approximation and sampling approximation with shrinkage to the gold standard permutation based approach. The p-values from our approach have strong correlation with the p-values from the gold standard method which improves after using shrinkage (Figure 3d). Critically, although this approach is obviously not exact, it provides an accurate approximation while being orders of magnitudes faster than standard permutation (Supplementary Note).

Finally we show how our approach can be used to identify sQTLs within sGenes. We show the FWER bound learned by our model for an example gene in Figure 3e. This bound corresponds to the critical values that controls FWER at a level equal to the BH threshold *α* = 0.022 for sGenes. For this dataset, this specific BH threshold controls FDR at a 0.05 rate. All tests in the gene with test statistics above this bound can be considered sQTLs and this procedure will maintain sGene FDR control at FDR = 0.05 and sQTL FWER control at level *α* = 0.022 under the null. Importantly, the lead SNP is guaranteed to have a test statistic above this bound if the sGene is significant and below if not significant. For comparison, we also show the unweighted max T bound and Bonferroni bound. The Bonferroni bound is too conservative and finds no sQTLs. The max T bound, while less conservative, misses sQTLs near splice sites that the weighted approach is able to detect. This example clearly illustrates the advantage of our approach for sQTL detection since we can analyze more than just the lead SNP which is prohibitively limited.

### MAJIQTL Refines Variant Prioritization and Effect Size Inference

Next, we evaluate the performance and utility of our effect size inference model and variant prioritization strategy. We begin with a simulation study to analyze the behavior of the betabinomial regression model for sQTL effect size estimation. Our aim is to understand whether the model is well calibrated for sQTL data and thus suitable for composite testing. We simulated splice junction and genotype data under 32 parametric conditions that can affect data factors such as MAF, skewness (*E*(Ψ)), overdispersion (Φ), coverage (*λ*), and changes in coverage (Y/N). The genotypes were simulated under Hardy-Weinberg equilibrium ratios based on the MAF. Further details are given in Supplementary Note. We evaluate the false positive rate and power of our model under the standard null hypothesis 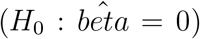 compared to two baselines: linear regression and linear regression with RINT transformed phenotypes (Figure 4a). We find that the Beta Binomial model is the uniformly most powerful and controls the false positive rate while the other models fail in some scenarios, as we detail next.

**Figure 4:**
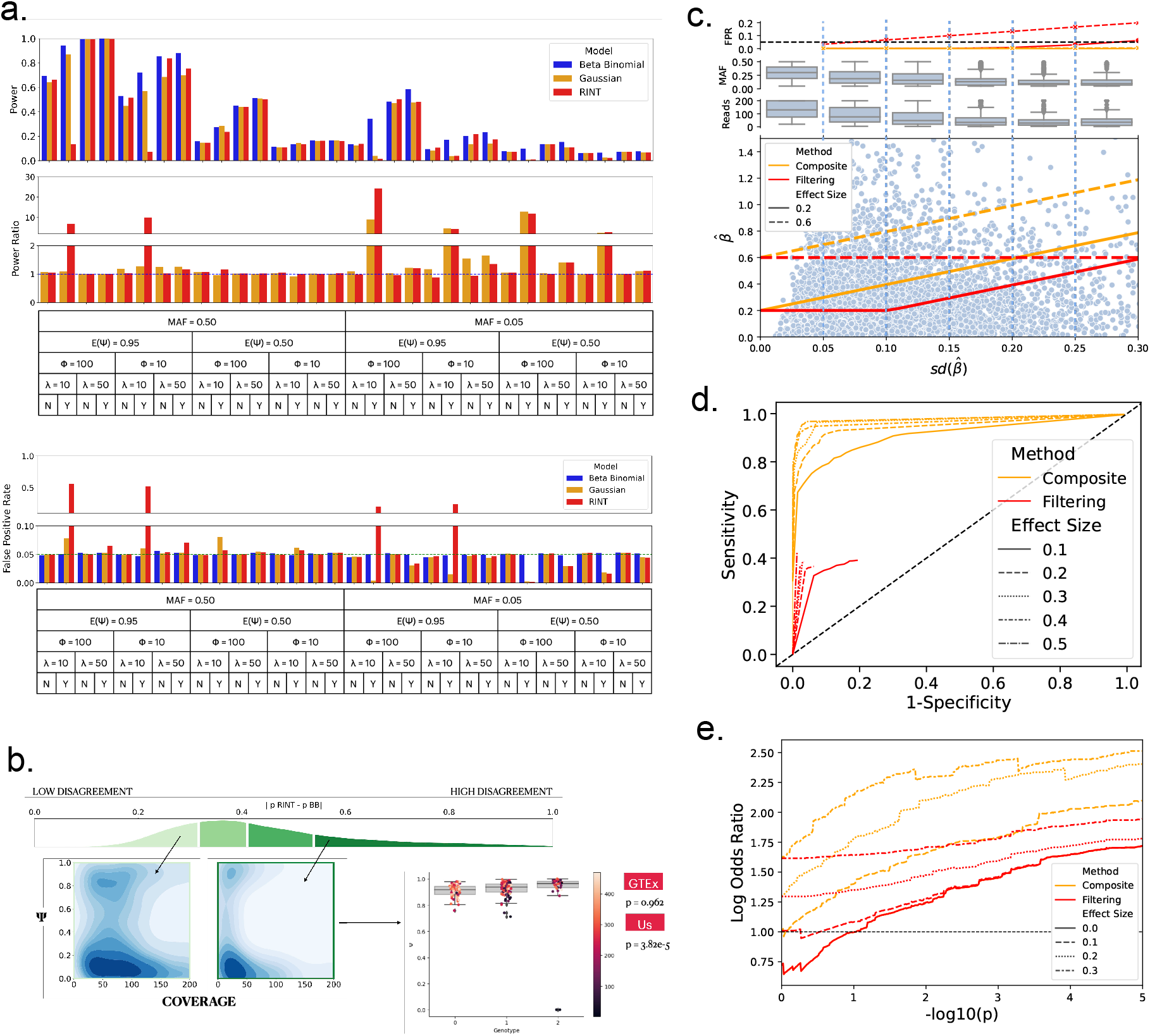
Evaluation of Composite Effect Size Inference. **(A)** Simulations comparing the power (top) and false positive rate (bottom) of the Beta-Binomial model, OLS regression and OLS regression with RINT transform across 32 parametric combinations for detecting sQTLs with non-zero effect size. The combinations are given by the chart on the x-axis. MAF is the minor allele frequency. *E*(Ψ) is the mean Ψ for the major allele. *ϕ* is the overdispersion. *λ* is the mean coverage in reads drawn from a Poisson distribution. *Y/N* refers to whether the genotype is correlated with coverage (i.e. is a cQTL). All simulations use an effect size of ΔΨ = 0.01. The cQTL coverage effect size is 20 reads. **(B)** A comparison of effect size estimates between Beta-Binomial and RINT OLS in brain - cerebellum. Effect sizes were divided into 4 bins based on the quantile of the differences between their p-values (green histogram). The models tend to disagree when coverage is low (< 50) and Ψ values are skewed to 0 or 1 (upper quantile). The models have much higher agreement when coverage is high (> 50) and Ψ values are near intermediate values (bottom quantile). The example boxplot shows a sQTL which has high disagreement between the models. However, the Beta-Binomial model is also robust to low coverage outliers and makes a more accurate effect size inference since GTEx is unable to call this sQTL due to the outliers. **(C)** A comparison of the decision boundaries of two approaches for constructing sQTL sets defined by a minimum effect size threshold: composite testing (orange) and filtering (red). For clarity, we only show decision boundaries corresponding to 0.05 FPR control for two thresholds: 0.2 (solid line) and 0.5 (dashed line). The composite approach has a non-linear boundary that increases with the standard error (x-axis) of the effect size estimator (y-axis). In contrast, the decision boundary of the filtering approach does not always account for the standard error. sQTLs with a Beta Binomial effect size estimator (blue points) greater than the decision boundary are considered to be in the minimum effect size set. The distribution of the MAF and read counts for sQTLs in each bin (blue dashed lines) on the x-axis (middle and bottom bar plots) indicates that these factors decrease when the variance increases. Composite testing also controls the FPR at the desire 0.05 level (albeit conservatively) while the filtering approach does not (top bar plot). **(D)** The ROC curve for the minimum effect size sQTL sets generated by the composite (orange) and filtering (red) approaches. Five effect size thresholds are shown from 0.1 and 0.5 and are represented by the various lines. The decision threshold for each approach is the sQTL p-value. For filtering, it eventually becomes impossible to separate the positive and negative classes by p-value since the effect size filter dominates, hence the red lines appear truncated. **(E)** Enrichment of minimum effect size sQTL sets for variants that disrupt splice sites (spliceAI Delta score > 0.2). The minimum effect size threshold for each set is indicated by the style of the line and the method used to generate the set is indicated by the color. The x-axis is the p-value (shown in -log10 space) rejection threshold for the method which can be interpreted as the FPR. The y-axis is the enrichment measured by the log-odds ratio. The composite and filtering approaches are equivalent at a threshold of 0 thus these two lines overlap.

The simulation results shown in Figure 4a offer important insights on the various modeling approaches behavior. First, as expected, RINT regression and OLS both fail to control false positives in the presence of heteroskedastic effects under the null. This is reflected in the simulations where the coverage is also associated with genotype and usually manifests in real data when a SNP is also an eQTL. We observed that the problem is also exacerbated at low MAF. Since OLS assumes homoskedasticity and uses a pooled variance model, the variance estimate is dominated by the variance of the largest genotype group. When allele frequencies are imbalanced, the variance can be systematically over or under estimated. In the case of RINT regression, the model also suffers from the assumption that the phenotype data is continuous. However, at low coverage, splicing quantifications can effectively be regarded as discrete. Specifically, when the phenotype in one genotype group is discrete (e.g. the total coverage is 10 and there are only 11 possible phenotype values) but continuous in another (e.g. the coverage is much higher), the ranks of the values cannot be compared. Doing so leads to bias in the mean estimate.

Another key point regarding effect size estimation that is illustrated by our simulation analysis in Figure 4a is the differences between fold changes and changes in ΔΨ. By design, the Beta Binomial model we use has substantially higher power to detect sQTLs with large differences in fold change instead of ΔΨ. In these simulations, we used ΔΨ as the effect size measure (set at ΔΨ = 0.01). However, when the minor allele mean is 0.5 and major allele mean is 0.52, ΔΨ is 0.02 but the change in the log odds ratio or Δ*log*(*Logit*(Ψ)) is 0.08. In contrast, when the minor allele mean is 0.95 and the major allele mean is 0.97, the effect size is still 0.02 in ΔΨ space but 0.53 in Δ*log*(*Logit*(Ψ)) space. Therefore, even through the effect size in terms of ΔΨ is constant in all simulations, the gain in power for Beta Binomial is larger when the minor allele mean is close to 0 or 1. We argue this property is desirable for two reasons: First, many disease associated phenotype occur with low MAF where the Beta Binomial gains more power. Second, previous work showed that alternative splicing can be modeled as competition between splice sites, each with its own usage rate [33]. Consequently, genetic variants that effect the splicing rate of a specific junction results in a non linear, sigmoid shaped, effect in Δ*Psi* space, depending on the initial rates ratio. For example, we may observe two isoforms that exist in a 1:1 ratio and thus have a Ψ of 0.5. A large 10 fold increase in 1 isoform caused by a variant would result in a 1:10 ratio. This corresponds to ΔΨ = 0.41 and Δ*log*(*Logit*(Ψ)) = 2.3. However, when the same 10 fold change is observed when the initial ratio is 1 : 10 and increases to 1 : 100, ΔΨ = 0.08 while Δ*log*(*Logit*(Ψ)) remains the same at 2.3. For this example, it can be argued that such a gain in power for ΔΨ = 0.08 may be marginal in some biological contexts and that confidently capturing such a change is hard. We note though that researchers can still filter hits by MAJIQ’s built in ΔΨ criteria and stress that under our model the significance of such a difference is still assessed given the observed coverage level and under composite testing (see below).

Next, we sought to assess the scenarios under which the Beta Binomial and RINT OLS tend to agree or disagree when analyzing real data. We divided the tests into 4 bins based on the absolute difference between the p-values of the RINT model and Beta-Binomial model. Then in each bin, we investigated the distribution of Ψ and coverage as shown in Figure 4b. The simulation study described above suggested the Beta-Binomial model has higher power to detect associations with large isoform fold change which occurs when the mean Ψ is close to 0 or 1. Inline with this, the results on real data shown in Figure 4b demonstrate the models tend to disagree the most when coverage is low and Ψ values are skewed to 0 or 1. When investigating cases of disagreement, we find that the Beta-Binomial model is robust to low coverage outliers (see example in Figure 4b).

Having shown the regression model is well calibrated for sQTL data, we then turn to evaluate the composite testing procedure. Without composite hypothesis testing, researchers will typically construct a rejection set using associations that are significant under the standard null hypothesis and filter sQTLs with observed effect size estimates below a desired threshold *θ*. It is instructive to first illustrate how composite testing differs from this baseline approach by showing the decision boundaries for both approaches (Figure 4c, bottom scatter plot). Composite testing produces a non-linear decision boundary [34]. Specifically, the threshold for the effect size estimator increases as its standard error increases thus more evidence is required to reject the hypothesis that the effect size is smaller than *θ* when the standard error is high. The primary contributors to higher variance in estimates include low junction read coverage and minor allele frequency (Figure 4c, top box plots). The filtering approach uses the standard null hypothesis decision boundary (equivalent to composite testing with *θ* = 0) but then applies a variance agnostic cutoff. This “mixture boundary” can completely ignore the variance of the estimator at high effect sizes (*θ* = 0.6 in the example). Figure 4c top graph demonstrates that as a result of the aforementioned differences in decision boundaries, composite testing controls the FPR at or below the desired level. In contrast, the filtering approach can exceed the desired FPR when the filtering component begins to dominate (see simulation details in Supplementary Note).

To evaluate the composite testing approach, we compare agreement between sQTLs in the composite rejection set and high confidence effect size annotations that we treat as ground truth labels. We cannot know the true effect size (i.e. population parameter) of each sQTL. Instead we use an information sharing strategy to combine estimates across tissues under the assumption that effect sizes are shared across tissues. This assumption has been used in many other studies [36]. Specifically, for a given sQTL, we compute the 95% confidence interval of its effect size estimate in each of the 5 representative tissues. Then we take the intersection of these intervals to be a high confidence range for the population parameter. To maintain a high quality set, sQTLs with tissue specific effect sizes in which confidence intervals across tissues do not all overlap are omitted. A sQTL is then considered to have an effect size greater than the desired threshold *θ* if the combined interval does not overlap and is not between *θ* and − *θ* (see Supplementary Notes for more details and illustrative examples). Using these high confidence effect size annotations as ground truth labels, we find that sQTL membership in the composite rejection set at varying FPR levels closely matches the labels as measured by AUC (Figure 4d). Importantly, for all thresholds *θ*, composite testing outperforms the naive filtering approach.

Finally, we show that using the composite sets to prioritize variants results in finding more sQTLs which are likely to disrupt splice sites usage, hence arguably more likely to have functional importance. For a given effect size threshold *θ*, we compute the rejection set at varying log10 p-value thresholds using either the composite p-values or original p-values. We then compute the enrichment of each of those rejection sets for splice site disrupting variants (Figure 4e). Here a variant was considered splice site disrupting if SpliceAI [37] predicted a change in splice site usage probability greater than 0.2. The 0.2 score is recommended by the authors for high recall. See Supplementary Note for further details of the enrichment calculation. We observe that the enrichment increases with effect size and p-value threshold. Importantly, the composite approach results in higher enrichment at all effect sizes.

### MAJIQTL Improves sQTL Enrichment in Functional Annotation and Identifies Novel sQTLs that Explain GWAS Signal for Neurodegenerative Disorders

Having shown the merit of each individual component of the MAJIQTL framework, we then use the full pipeline to call sGenes and analyze the functional enrichment of their sQTLs. Specifically, we use the weighted multiple testing method to call sGenes and call sQTLs that pass the weighted FWER bound for each sGene controlled at the BH threshold. We compared our approach to the std-sQTL approach which uses the max T procedure (unweighted FWER) to discover sGenes with Leafcutter quantifications and selected sQTLs within sGenes that passed a gene level Bonferroni threshold. We analyzed enrichment of the sQTLs for two SNP functional annotations: splice site disrupting variants and RBP binding sites. A sQTL was considered splice site disrupting if it was predicted to have a SpliceAI delta score greater than 0.2 for the 5’ or 3’ splice site of the sQTL’s splice junction. We considered a RBP to bind to a sQTL if it had an eCLIP peak that overlap the variant in the ENCODE dataset for any of the 113 RBPs in K562 cell lines. Binding was determined using an IDR threshold which is the stringent cutoff recommended by ENCODE. We find that sQTLs in MAJIQTL sGenes are enriched for these annotations compared to sQTLs in non sGenes at a 0.05 FDR level across all 5 tissues based on Fisher’s exact test (Figure 5a). The std-sQTL pipeline using Leafcutter has lower enrichment in all tissues. Notably, we found that only using lead SNP approach did not yield sufficient enrichment for either method due to the highly varied positions of lead SNPs which may not be casual.

**Figure 5:**
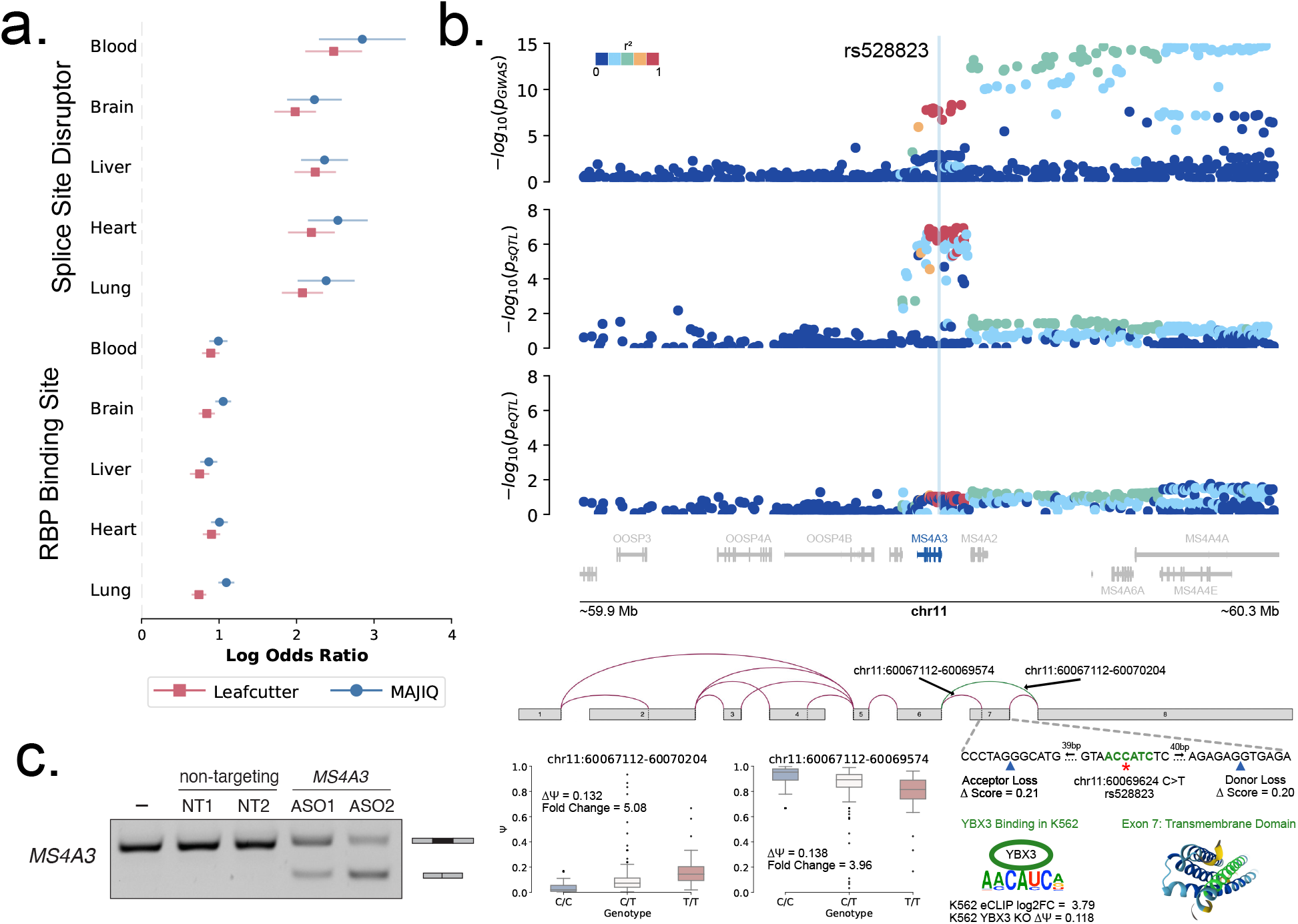
Functional Analysis of sQTLs. **(A)** Enrichment of sQTLs proximal to splice sites within sGenes discovered by MAJIQTL (blue) for two functional annotations: splice site disruption and RBP binding. Leafcutter (red) sQTLs are found using the std-SQTL pipeline and the Bonferroni threshold in each sGene. A variant is considered splice site disrupting if it has a SpliceAI Delta score > 0.2. A variant is considered a RBP binding site if binding of at least one of one RBP is observed in the ENCODE eCLIP data (K562). The x-axis shows the log-odds ratio for enrichment and the y-axis shows the tissues. The confidence interval is the 95% interval for the log odds ratio computed using Fisher’s exact method. **(B)** Analysis of the variant rs528823 (vertical blue line) and its role in disease. The Manhattan plots show the Alzheimer’s GWAS (top), MS4A3 exon 7 junction sQTL (middle) and MS4A3 expression eQTL (bottom) p-values for all variants in a locus around the gene (highlighted in blue) on chromosome 11. The QTL tissue is blood. The points are colored by their LD with the variant of interest. The splice graph shows all junctions (green denovo, red annotated) in the gene. The junctions of interest are marked by the black arrows. The variant rs528823 is located on exon 7 at the position indicated by the red star. The effect size of this variant on the marked junctions is shown in the box plots (bottom left). The variant has > 0.2 spliceAI delta score at the indicated splice sites (blue arrow). The bases highlighted in green show the position of the YBX3 binding motif. YBX3 binds at this location based on ENCODE eCLIP data and alters splicing based on the splicing change observed in YBX3 KD. Exon 7 encodes a transmembrane domain of the protein. **(C)** PCR reactions in K562 cell lines visualized on a agarose gel for the MS4A3 splicing event. From left to right there are 5 conditions: control (no ASO blocking), two ASOs with non-targeting control sequences (Supplementary Note), and two ASOs that targeting exon 7 of MS4A3 (Supplementary Note). When the ASO successfully blocks the splice site on exon 7, we see isoforms where exon 7 is skipped (smaller fragment further down the gel).

Finally, we analyze the sQTLs discovered by the MAJIQTL pipeline to understand if they can provide new insight on disease mechanisms. We first obtained summary statistics for Alzheimer’s and Parkinson’s disease from the studies Kunkle et al. 2019 and Nalls et al. 2019 respectively. We find that the sQTLs identified by MAJIQTL explain 11% and 8% more GWAS variants for Alzheimer’s and Parkinson’s respectively compared to std-sQTL (Supplementary Note). Using our variant prioritization approach, we identify rs582283 in the composite rejection set at a 0.2 composite level as a variant of interest which may be implicated in disease (Figure 5b). This variant is associated with the splicing of exon 7 in the gene MS4A3 (p=3.55e-7) but does not appear in the GTEx catalogue as a sQTL for this splicing event. It is also a GWAS signal for Alzheimer’s (p=1.27e-8) and a significant sQTL in blood but surprisingly not an eQTL (p=5.51e-2). The variant is observed to have strong LD with other variants in the gene but not variants outside of the gene. Since the variant is not significant for expression, this points to splicing disregulation as a potential mechanism underlying the disease association. Inline with this hypothesis, we find rs582283 is located on exon 7 which is in a cassette event with a denovo junction. Exon 7 is an important transmembrane domain and exclusion may have implications for loss of protein function. The variant appears to promote skipping of exon 7 with an estimated effect size of ΔΨ = 0.132 or 5.08 fold when the variant changes from a C to T. Furthermore, the variant is predicted to be a splice site disrupting variant with a SpliceAI score change for each adjacent splice site (5’ and 3’) of 0.2. The variant overlaps the binding motif for the YBX3 RNA Binding Protein and the position of the variant in the motif is non degenerate suggesting that the change in variant significantly reduces binding affinity. Supporting this hypothesis of rs582283 disrupting RBP binding, when we target this region with an antisense oligonucleotide (ASO) we observe a significant change in splicing (Figure 5c). Details for the experiment are describe in Supplementary Note. Furthermore, ENCODE K562 eCLIP experiments suggest YBX3 binds to this position (Log2FC = 3.79 increase compared to background), while shRNA knockdown of this RBP reduces the inclusion rate of the exon compared to controls (ΔΨ = 0.118).

## Discussion

In this work, we decompose the current state of the art sQTL analysis pipelines into their fundamental components and individually evaluate each component on five representative GTEx tissues. The results of this case study set a precedent for future work by identifying three key limitations with existing pipelines and demonstrating how they could be addressed to improve sQTL discovery and analysis. First, we show that using the Leafcutter splicing representation can miss crucial splicing events like intron retention and alternate transcript starts/ends which represent a significant number of sQTLs that explain GWAS signal. This result is inline with other works demonstrating the importance of capturing additional types splicing variations. For example, our previous work experimentally verified the effect of rs6410, a variant associated with skeletal aging, in promoting retention of intron 3 of CYP11B1 [38]. More generally, several other studies have highlighted the importance of integrated splicing representations, finding more variants explaining disease loci through full isoform and alternate transcript end (APA) QTLs [18, 39, 40].

Second, we show that the current statistical methods used for sGene/sQTL discovery are under powered and not well calibrated. It has been suggested that QTL discoveries are saturated and improvements to statistical techniques are unlikely to further close the colocalization gap [41, 42]. However, our results indicate that using multiple hypothesis testing correction based on FWER control for sGene discovery is conservative compared to alternative methods. This conclusion is concordant with the findings of a recent study that benchmarked methods for isoform QTL discovery [43]. We show that improved statistical modeling finds new sGenes harboring sQTLs which are enriched for various functional annotations and variants implicated in neurodegenerative disease. In addition, through comparison of methods for sQTL effect size inference, we show that the heteroskedastic and discrete characteristics of splicing data necessitate the use of a Beta-Binomial model. Importantly, the effect sizes of sQTLs that are also eQTLs cannot be reliably estimated by OLS due to inconsistent variance in splicing quantifications induced by differences in gene expression and coverage. The resulting inflation of false positive non-zero effects serves to illustrate that simply more hits is not always better, and carefully assessing the models’ fit to the data is critical. In a broader context, we suspect that this model misspecification is partially responsbile for the widespread inconsistent reporting of overlap between eQTLs and sQTLs [4, 18, 21].

Third, we illustrate the limitations of current practice for SNP and junction selection in cis sQTL analysis. This topic is seldom discussed in recent QTL work. Instead, most sQTL studies opt to borrow the selection parameters directly from eQTL studies, using the same large window size (1Mb), discarding splicing events with even 10% missing values and imputing the rest [21, 28, 29]. We show this current standard for study design can severely decrease detection power by needlessly increasing the multiple hypothesis testing burden and potentially filter out junctions with missing values that may harbor strong sQTL signals. However, we show this problem can be easily mitigated by utilizing power informative covariates (splice site distance, missingness rate), emphasizing the importance of not discarding data.

To address the issues from this cases study, we developed here MAJIQTL, a novel sQTL discovery toolkit with two new statistical methods to improve sQTL mapping and facilitate downstream analysis. First, we introduce a weighted multiple hypothesis testing method (which can be seen as a sGene test) that leverages SNP and junction level covariates to significantly increase sGene detection power. We note such weighted control procedures are not new, but previous methods have not been adapted in the QTL field due to their focus on FDR control and broad applicability [19, 44, 45]. The appeal of our method lies in its design tailored for QTL studies which requires a FWER based approach for false positive control in the presence of complex LD and junction correlation structure [30, 46]. Beyond increased power, we show that controlling the FWER of each sGene at the BH critical value allows for individual sQTLs to be called at a constant FWER while maintaining gene level FDR control. Despite the widespread use of lead-SNP analysis due to current methodological limitations [18, 40, 47], we highlight how our approach which uses multiple sQTLs per gene leads to increased enrichment for splicing relevant functional annotations.

Second, we propose using a composite Beta-Binomial model for effect size inference and variant prioritization. Unlike previous applications of similar models for sQTL detection [15], we focus on the utility of using the model parameter as an interpretable measure of sQTL effect size. Notably, there is currently no consensus definition for sQTL effect sizes reported on the GTEx portal despite extensive use of the allelic fold change metric for eQTLs [32]. For RNA splicing, two common measures are fold change and ΔΨ. The latter is arguably the most commonly used unit for studying splicing regulation though several studies have advocated for using fold change to represent the effect of genetic variants on splicing [33, 48]. In our case, the fold change arises as the natural effect size from the Beta-Binomial model, though we note that any identified sQTL can also be filtered by MAJIQ’s ΔΨ as well. We show that the fold change effect size estimates of our model are both accurate and informative of functional annotation. We also use a composite testing method to create minimum effect size sets as a way to prioritize variants. A related approach using fine mapping showed that variants in fine mapped credible sets for QTLs are enriched for disease heritability [49]. Similarly, we show our composite sets are enriched for important functional annotations such as splice site disruption and RBP binding. This provides a complementary set of annotations which we demonstrate are useful for variant prioritization. We show that the naive approach of selecting significant sQTLs (with non-zero effect size) followed by filtering for a desired effect size results in lower accuracy and variant enrichment. Although we only investigate the benefits of minimum effect size sets, we emphasize that composite testing can also be used to create maximum effect size sets. Identifying such sets may have potential application for developing therapeutics since variants exerting small effect sizes on splicing with a strong phenotype are more likely to be efficiently blocked by antisense oligos (ASO).

Beyond sQTL, we believe our effect size model and annotations can be used for other applications that involve predicting genetic effects on splicing. For example, splicing TWAS is an alternative approach for identifying risk variants that are mediated by splicing. However, current models still rely heavily on multivariate linear regression for imputing splicing phenotypes [50]. It is not clear whether focusing on a multivariate strategy is better than a parametric strategy since our results suggests that using the correct error model is crucial for estimating the genetic component of splicing effect size. Our results can also be used to train and validate deep learning models that predict tissue specific splicing from DNA sequence. Similar ideas have been used for expression prediction models with eQTLs [51]. It is generally believed that cis-QTLs are shared across tissues but may have tissue specific effect sizes [36]. We identified many instances of tissue specific effect sizes by using composite testing and confidence intervals which can be used to train tissue specific prediction models. Although several methods have been developed to predict tissue specific Ψ from genomic sequence [52–55], the most successful methods for predicting the effect of genetic variants can still only predict splice site location which is effect size and tissue agnostic [37]. This work thus has the potential to lead to improvements in tissue specific splicing code models.

Finally, through applications to real GTEx data, we show that our method discovers many novel sQTLs which are also significant in Alzheimer’s and Parkinson’s GWAS. We highlight a new variant, rs528823, which may be linked to Alzheimer’s through splicing rather than gene expression. Specifically, the variant appears to disrupt the inclusion of an exon in the gene MS4A3 by perturbing a YBX3 binding motif, a hypothesis supported by our ASO targeting of this site as well as YBK3 CLIP binding and shRNA KD. Although our result is only significant in blood, other genes in the MS4A gene cluster have been previously implicated in Alzheimers through immune cell related functions which may act through blood [56].

There are several limitations with our approach that are important to highlight. First, isoform quantification from short read data are generally considered to be inaccurate. There is no guarantee that isoform level sQTLs discovered are real due to errors in the quantification. However, at the pace that long read technology is improving, we believe these quantifications can be easily replaced with long read data and our approach supports that substitution. Second, we don’t address trans-QTLs in this work. While MAJIQTL can recover many common and cis variants associated with splicing, many variants that colocalize with GWAS signal can be rare and trans-acting. Detection of these variants requires complementary approaches. Specifically, splicing code models can reveal rare variant effect on splicing using a single patient’s DNA sequence. Such models have been effective in finding cryptic splice site resulting from variants with low frequency in populations [57]. Trans-QTLs are generally mediated by cis effects. For example, a distal regulatory splice factor could have cis QTLs which are trans QTLs for distal splicing targets. Modern approaches have used CRISPR or Perturb-Seq screens to knock out regulatory factors and associate cis-QTLs of these factors with splicing of their target genes[58]. Finally, our study is focused on methodological advancement rather than exhaustive and robust sQTL mapping. As such, it does not include a replication study. Assessing replication is an important component which helps land confidence in both the methods and consequent results. However, we caution that replication alone does not necessarily indicate correctness. Specifically, several of our results point to the issue that misspecified models can consistently produce recurrent false hits.

In conclusion, we believe that MAJIQTL will significantly advance sQTL discovery given the many improvements it offers over existing methods. Furthermore, while the pipeline integrates with the MAJIQ tool set, it is compatible with existing tools such as Leafcutter, rMATs and QTLTools. We hope the genetics community will take full advantage of this new method for RNA splicing analysis and potentially modeling of other QTL.

## Methods

### A Weighted FWER Control Model for sGene Discovery

Here, we present a method for cis sGene discovery that leverages splice junction and SNP covariates to improve detection power. For a gene *g* ∈ [*G*] with *J*_*g*_ splice junctions and *K*_*g*_ SNPs in a predefined genomic window around the gene, consider the set of *J*_*g*_ *×K*_*g*_ marginal hypothesis 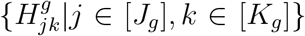 with corresponding test statistics 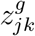. We assume that these test statistics are computed using linear regression between each pair of junctions and SNPs but do not assume they are independent under the null. Let a sGene be defined as a gene with at least one true alternative marginal hypothesis. Then to test for sGenes in the setting without covariates, we wish to reject the complete null hypothesis *H*_0_. Formally,

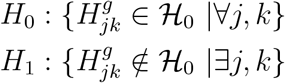

This hypothesis can be tested for each gene using 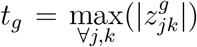 as the test statistic, as described in Westfall and Young 1993 [59], to compute an unweighted gene level p-value. The p-value is given as

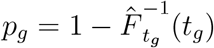

where 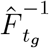 is the CDF of the null distribution of *t*_g_ which is typically estimated using a permutation based approach (we describe the details in a later section). Then a false discovery rate (FDR) control procedure such as Benjamini Hochberg (BH) is applied to control the sGene FDR at a desired level across all genes by determining the sGene p-value decision threshold *α*. This approach for genome-wide FDR control is equivalent to controlling the family-wise error rate (FWER) for each gene at level *α*. Specifically,

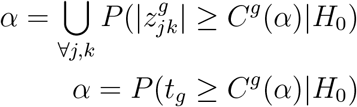

where *C*^*g*^(*α*) is the uniform critical value for FWER *α*. We emphasize that *C*^*g*^(*α*) is different for each gene. Now consider the setting where we have covariates 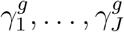 for each junction and 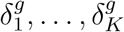 for each SNP in gene *g*. In this study, we treat the junction covariates as the missing value rate *γ* ∈ [0, 1] and the SNP covariates as the normalized proximity to the splice site which is normalized w.r.t to the maximum allow window size in bases *δ*∈ [0, 1]. Critically, these covariates must be independent of the p-value under the null [60]. However, if the covariates are correlated to 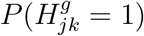, we can specify weighted critical values 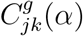 that allocates more of the FWER budget to tests where the marginal alternative hypothesis is more likely to be true subject to the constraint that the FWER is still controlled at exact rate *α*. To derive these weighted critical values, we factor 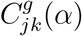 into a covariate dependent weight component *w*_*jk*_ and a gene level constant Δ_*g*_ (*α*) dependent on *α* that is used to maintain the FWER budget.

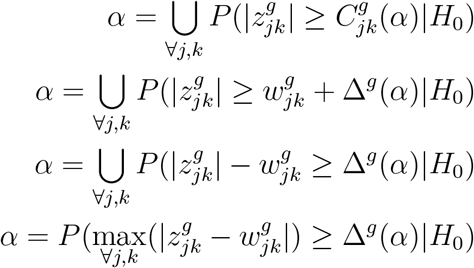

The proof for this decomposition is given in Supplementary Note X. Then, in contrast to the unweighted p-value, the weighted p-value per gene can be computed as

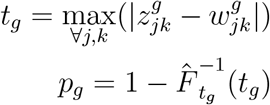

Assuming the weights 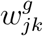 are known, the weighted FWER rejection bound 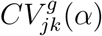 can then be recovered by choosing Δ_*g*_ (*α*) such that

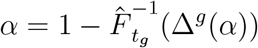

Individual sQTLs within sGenes can be identified if 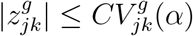.

In the remaining sections, we discuss 1) how to estimate 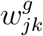 to increase the number of sGene discoveries and 2) how to estimate 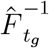. It is important to emphasize that the FWER under the complete null is controlled for this procedure for any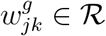, even if 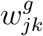 is not chosen optimally.

### Weight Optimization

Now we describe a fast approach for choosing the weights 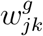 that increases the number of sGene discoveries. This method provides an approximately optimal solution that scales to hundreds of millions of tests. Our goal is to learn some function *g* that maps covariates 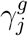 and 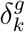 to their corresponding weight 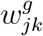 such that the number of sGene discoveries is maximized. It has been shown that the theoretical optimal quantity for the weights is the probability of the alternative hypothesis 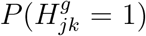. This is often called the local false discovery rate (lFDR) [61]. Consider the two group model for the observed test statistics as described in Efron 2001 [62]. Under this model, we assume that the test statistics follow a mixture of two normal distributions. One mixture component represents the null distribution *f*_0_(*z*) while the other represents the alternative distribution *f*_1_(*z*). In the setting with covariates, the mixing proportions *π* and alternative distribution depend on the covariates. Formally,

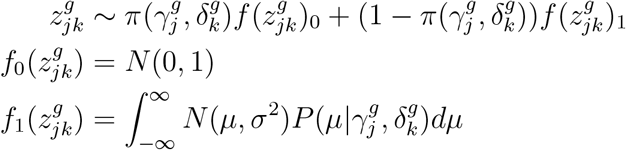

The lFDR can then be formulated as the posterior probability of the alternative hypothesis conditioned on the covariates.

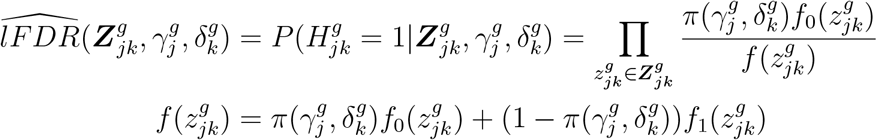

If we could successfully perform inference for the above model, we would have our mapping function *f*_*w*_ and could stop here. However, maximum likelihood inference of the model parameters is challenging and generally not feasible at scale [44, 62]. Instead, tractable solutions often use a data driven approach to derive weights. In brief, these methods search over the space of all possible weights subject to a budget constraint to find the values that maximizes hypothesis rejections in the dataset. Although this objective appears to be NP hard, various approximations have been proposed [19, 45]. We use a simple 3 step procedure to approximate the optimal weights conditioned on covariates. We outline the steps and advantages of this approach below.

1. Compute the pseudo lFDR estimator 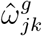 which has the property 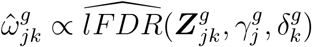. This is a cheap plug-in estimator that does not require numerical optimization.

2. Learn the function *g*(*γ, δ*) that maps the covariates 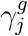 and 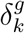 to 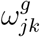 using a Guassian Process (GP) regression model trained on the estimators computed in (1).

3. Estimate the global scaling factor *λ* such that *w*_*jk*_ = *h*(*λω*_*jk*_) maximizes the number of sGenes discoveries for some function *h*.

In step 1, we describe how to compute 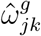. We propose using the Kolmogorov-Smirnov (KS) distance 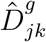 between *f*_0_ and *f* as the estimator for 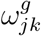. The primary advantage of this approach is that it does not require any numeric optimization. Following Efron 2001, it can be shown that 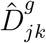 is proportional to the upper bound on the maximum likelihood estimate for lFDR and is thus a reasonable choice of estimator. We assume that the density of *f* (*z*) (the observed mixture distribution of test statistics) is typically smooth and approximately follows a normal distribution with a heavy tail. This is supported by empirical evidence across multiple studies, including our own [62, 63]. Given this approximation for *f* (*z*),

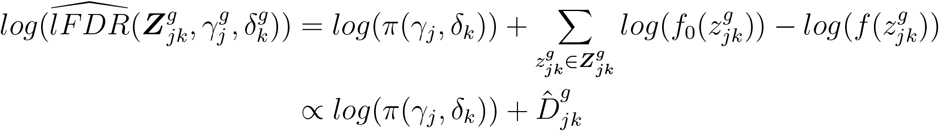

In other words, 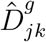 is proportional to the the log likelihood ratio of the null distribution and observed test statistic mixture distribution. We can consider *π*(*γ*_*j*_, *δ*_*k*_) as a nuisance parameter since the ratio 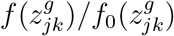 provides the upper bound on *π*(*γ*_*j*_, *δ*_*k*_). Specifically, given the approximate density of 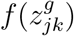,

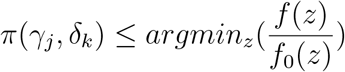

since *π*(*γ*_*j*_, *δ*_*k*_) must satisfy the constraints 0 ≤ *π*(*γ*_*j*_, *δ*_*k*_) ≤ 1 and 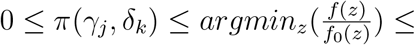 1. Thus it remains that when

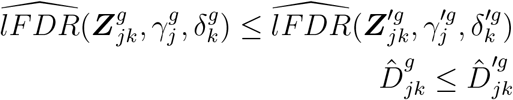

since 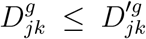 implies the upper bound of 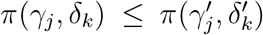. Here, the prime (‘) notation indicates any other subscript *j* and *k*.

To compute, 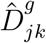, we use a pooling strategy across genes and nearby covariates. For a local pooling area *ν* = 0.025, let 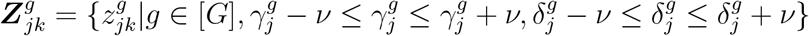. The empirical CDF 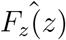 and 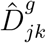 can then be computed as

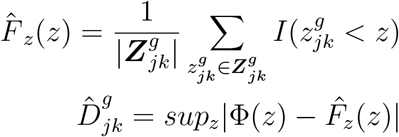

where Φ is the standard normal CDF. This gives us the the estimator under the assumptions that the lFDR conditioned on the covariates is similar across genes and nearby covariates. Then we can obtain bootstrap samples for the estimator by sampling from 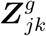 with replacement and computing 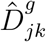 on the bootstrapped samples.

In step 2, we learn a function mapping from covaraties 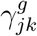 and 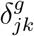 to 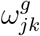. We model this function *g* with a Gaussian Process (GP) prior. The normal error model is used to approximate the the variance of the estimator across genes.

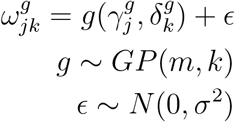

Here, *m* is the zero mean function and *k* is the RBF kernel. To train the model, we generate training datasets 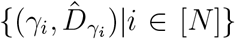 and 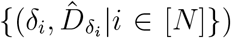 using the pooling strategy described in (1) for up to *N* samples. For *γ*_*i*_ and *δ*_*i*_, we sample covariates uniformly from [0,1]. Then for the given *γ*_*i*_, generate generate set 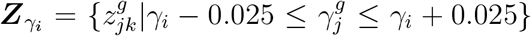 and 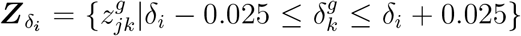. We then bootstrap estimators 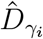 and 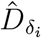 from sets 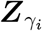 and 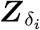 to form covariate pairs 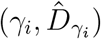 and 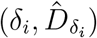. Then we train the model on these datasets using 10 fold cross validation to avoid over fitting. For inference, we use the mean of the posterior predictive distribution as the weight. Optimization is preformed using the GP regression function in scikit-learn [64].

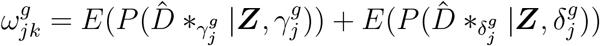

In step 3, once we have predictions for 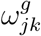, we can then perform optimization of *λ* to maximize sGene discoveries. This is obtained by choosing *λ* such that for some FDR threshold *α*

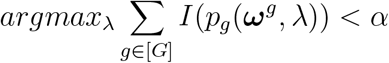

Here, *p*_*g*_(***ω***^*g*^, *λ*) is the weighted p-value of gene *g* computed using 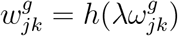. We choose *h* to be the exponential function which empirically produced the best results on our datasets.

### Estimation of the Null Distribution

Here we show how to estimate the CDF of the null distribution 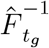. We first describe an approach for sampling from this null distribution wtihout using permutations. This approach can decrease the run time by an order of magnitude for population level studies and can be used with or without weighting.

Assume the setting where we have *J* splice junctions and a single fixed SNP *k*. We perform *J* association tests and obtain the test statistic vector ***z***_*k*_ = (*z*_1*k*_, *z*_2*k*_, …, *z*_*Jk*_). Under the null, ***z***_*k*_ ∼ *MV N* (**0**, Σ_*zk*_) and the marginals *z*_*jk*_ ∼ *N* (0, 1). Similarly, considerthe setting where we have *K* SNPs and a fixed splice junction *j*. In this setting ***z***_*j*_ = (*z*_*j*1_, *z*_*j*2_, …, *z*_*jK*_) and ***z***_*j*_ ∼ *MV N* (**0**, Σ_*zj*_). Taken together, we can sample the null distribution of these test statistics from a matrix normal distribution *MV N* (**0**_*J*_*×*_*K*_, Σ_*zk*_, Σ_*zj*_). A sensible approach to estimation is to use the standardized sample covariance estimator (i.e. correlation) of the junction quantifications 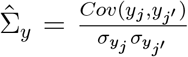 and genotypes 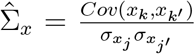 to approximate Σ_*zj*_ and Σ_*zk*_ respectively. It can be proven than Σ_*y*_ = Σ_*zk*_ or Σ_*x*_ = Σ_*zj*_ for OLS (Supplementary Note). With the parameters of the matrix normal estimated, we can then sample from this distribution *N* times and calculate 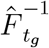 as follows

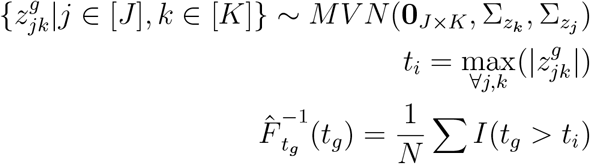

Briefly, sampling from a multivariate normal requires an affine transform of random independent standard normals *Z*. Specifically:

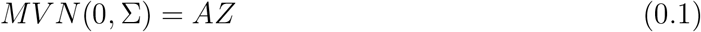

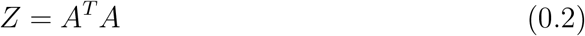

The matrix A is a symmetric matrix found using Cholesky decomposition. However, when Choloesky fails due to a matrix not being full rank, we resort to SVD to obtain decomposition *USV* such that 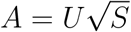.

To improve the accuracy of the covariance estimators, we use two adjustments. First, since it is often the case that *K >> N*, we use the Ledoit Wolf shrinkage estimator [65] instead of sample covariance estimator 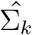 which is more accurate in this scenario. Specifically, it can be shown that the Frobenius norm between this LW estimator and population covariance matrix is minimized.

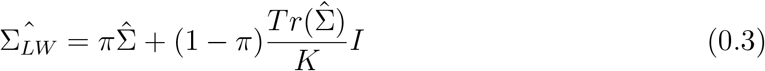

### Effect Size Inference Model for sQTL sQTL Model

For a splice junction *j* ∈ [1 … *J*] in sample *i* ∈ 1 … *M*], let *y*_*i*_ be the number of reads mapped to that splice junction and *n*_*i*_ be the total number of reads mapped to the splicing event (LSV, intron cluster, etc). For each sample *i*, we also observe genotype *x*_*i*1_ ∈ [0, 1, 2] which is encoded as the minor allele count. We also have *K* −2 covariates *x*_*i*2…*i*(*K*−1)_ and the model intercept *x*_*iK*_ = 1. The covariates often include known confounding factors that affect the genotype (e.g. population structure) and splicing data (e.g. RNA-seq batch) and unknown confounding factors which are inferred using an external model (e.g. MOCASSAIN[66], PEERS[67]). The association between the genotype and splice junction in each sample is modeled as a beta binomial regression. Note that our model is not a GLM (and thus does not benefit from generic solutions for GLMs) because the beta binomial distribution is not in the exponential family. The model likelihood is defined as

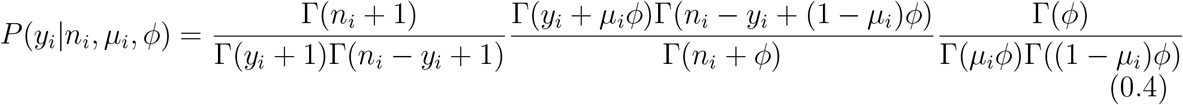

Here, *µ*_*i*_ is the mean of the distribution and *ϕ* is the dispersion. The gamma function is denoted as Γ(.). We choose a logit link function for our model. Thus *µ*_*i*_ is a linear function of *x*_*i*_ transformed by the inverse logit or logistic function such that

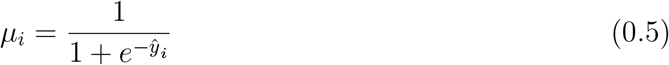

and

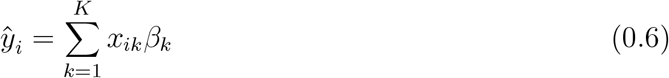

where *β*_*k*_ is the *k*_*th*_ regression coefficient.

To compute the p-value for *β*_1_ (recall that the index 1 refers to the genotype index), we use Wald’s method. The Wald test statistic and asymptotic distribution is given by

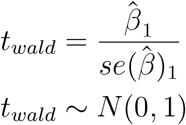

We compute the standard error as the following where *H* is the Hessian matrix.

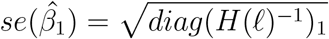

Note that our computation of the Hessian when evaluated at the MLE parameters produces standard errors that are more precise than the approximate Hessian calculated by black box solvers (Supplementary Note).

### Composite Testing for sQTL Effect Size

For composite testing, we follow the approach of Love et al. 2016. The null and alternative hypothesis are given as follows.

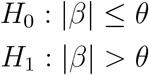

Let *θ* be the desired effect size threshold. Then the composite p-value is given by

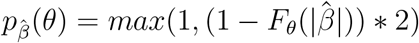

where *F*_*θ*_ is the CDF of 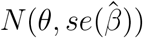. When *θ* = 0, this reduces to the null hypothesis *H*_0_ : *β* = 0.

## Data and code availability

Code and data will be released upon publication.

## Acknowledgements

We would like to acknowledge Dr. Caleb Radens, Dr. Iain Mathesion and Dr. Bogdan Pasaniuc for helpful discussion and advice. CDB passed away during the work on this project. We miss him dearly and believe he would have been proud of the final product. We dedicate this work to his memory and contributions to science.

## Author Contributions

DW and YB conceived the project. DW and YB developed the methods and planned the experiments with input from MG and CB. DW wrote the code and carried out the experiments and analysis. SJ handled all aspects of the software related to MAJIQ integration, including developing the visualization tools. DW and YB wrote the manuscript. All authors read and approved the final manuscript.

